# Flower clades and fruit clades: trade-offs in color diversification across angiosperms

**DOI:** 10.1101/2025.06.20.660771

**Authors:** Miranda Sinnott-Armstrong, Leah Maier, Stacey D. Smith, Agnes S. Dellinger

**Affiliations:** Department of Biological Sciences, Purdue University, West Lafayette, IN USA; Department of Ecology and Evolutionary Biology, University of Colorado-Boulder, Boulder, CO USA; Department of Botany and Biodiversity Research, University of Vienna, Rennweg 14, Vienna, 1030, Austria

**Keywords:** Angiosperms, evolutionary lability, flower color, fruit color, pollination, seed dispersal, stochastic character mapping, trait diversification, transition rates

## Abstract

**PREMISE:** Flowers and fruits are two major phases of plant reproduction which often use colorful signals to attract animal mutualists. Fleshy fruits develop from the ovaries of flowers, and both organs use the same suites of pigments to create color. Due to these developmental links and functional similarities, we sought to test for correlations in flower and fruit color lability across clades.

**METHODS:** We selected 51 clades (2989 species) of animal-pollinated and animal-dispersed plants and scored flower and fruit color into eight discrete (human-perceived) categories for the same set of species in each clade. We used stochastic character mapping to estimate the number and rates of transitions among colors in flowers and fruits.

**KEY RESULTS:** We demonstrate a negative correlation in the number of transitions in flower and fruit color across clades (R^2^ = 0.48, p < 0.001). Among animal-pollinated and animal-dispersed clades, the majority (67%) are “fruit clades” biased towards fruit color lability, while a minority (29%) are “flower clades” biased towards flower color lability. We further show that clades with many yellow- or orange-flowered species also tend to have those colors in their fruits, and that red flowers are more common in “flower clades” while green and brown fruits are more common in “fruit clades”.

**CONCLUSIONS:** These patterns suggest that clades specialize on one phase of reproduction or the other. Possible explanations include constraints on energetic investment into either pollination or dispersal, and environmental factors that select for color diversification in one organ but not the other. This variation is also likely shaped by the underlying structure of pigment pathways that contribute to color in both organs.

## INTRODUCTION

During the life cycle of flowering plants, the ovary at the center of the flower matures into the fruit. Many plants produce animal-pollinated flowers and/or animal-dispersed fruits that use signaling traits to attract their mutualistic partners (Waser et al. 1996; Van Der Pijl 2012). Despite the developmental continuity and functional similarity between these two organs, flowers and fruits are usually studied as separately evolving entities. Cross-organ correlations have been better studied between floral and vegetative traits and have been detected in various systems (Berg 1960; Armbruster et al. 1999; Brock and Weinig 2007; Meng et al. 2008). Similar assessments of flower and fruit traits, critical for understanding the evolution of plant reproductive systems, are scarce (Primack 1987; but see Kang and Primack 1991). At the genetic level, the morphology of reproductive organs is often shaped by highly pleiotropic loci (Armbruster 2002; Hall, Basten, and Willis 2006), and species with larger flowers tend to have larger fruits (Griffiths and Lawes 2006) and larger fruits in turn may have more (Bawa et al. 2019) and/or larger seeds (Sakai and Sakai 1995). These patterns hint at the possibility of correlations in other traits (e.g., color, volatiles) between flowers and fruits. However, the historic disciplinary focus on either flowers or fruits has limited our understanding of the evolutionary (in)dependence of these organs, thereby restricting our understanding of the entire reproductive pathway of flowering plants.

Color is one of the most important signaling traits in flowers and fruits, functioning in the attraction of animal mutualists. Flowers and fruits generally rely on the same biosynthetic pathways for producing pigments, primarily anthocyanins, carotenoids, chlorophylls, and betalains (Tanaka, Sasaki, and Ohmiya 2008; Grotewold 2006; Ranganath 2022), which give rise to the majority of plant colors (except for a few lineages with structural colors; e.g., Lee 1991; Middleton et al. 2020; Sinnott-Armstrong et al. 2022; Moyroud et al. 2017). Animal pollinators and dispersers have been identified as major selective agents of both flower and fruit colors, contributing to trait diversity in both phases of angiosperm reproduction primarily through shifts in pigmentation (Wheelwright and Orians 1982; Valenta, Nevo, and Chapman 2018; Schiestl and Johnson 2013; Stankowski and Streisfeld 2015) or other mechanisms of color production (e.g., addition of structural color to pigmented fruits; Sinnott-Armstrong et al. 2023). The developmental continuity, pleiotropic linkages, reliance on the same biochemical toolkit, and functional similarity (attracting animals) all suggest that flower and fruit colors may influence each other’s evolution.

At the same time, other factors suggest that flower and fruit colors evolve independently. Despite the similarities in the types of pigments used, the regulation of pigment pathways can be tissue-specific (Jaakola 2013; Muhlemann, Klempien, and Dudareva 2014; but see Berardi et al. 2016), allowing flowers and fruits to produce different pigments at different intensities even within an individual. Accordingly, surveys at regional scales indicate distinct patterns of flower and fruit color diversity, with differences both in the range of colors present as well as their prevalence (for instance, white is much more common in flowers while black and red are much more common in fruits; Wang et al. 2022; Delmas, Kooyman, and Rossetto 2020; Sinnott-Armstrong et al. 2018; Dyer et al. 2020). Moreover, pollinators and seed dispersers differ in their visual abilities, which results in different selection pressures that may lead to divergence in color signals to match the different sensory preferences of pollinators (mostly insects) and dispersers (mostly birds and mammals; Wheelwright and Orians 1982; Schiestl and Johnson 2013). These macroecological patterns suggest some degree of decoupling between flower and fruit colors, allowing their diversity to be shaped by different selective factors (e.g., Dellinger et al., 2025). These factors suggest that flower and fruit colors may evolve independently, even with their developmental connections and overlapping biochemical capabilities.

Given the fundamental role of color in flower and fruit ecology as well as the extensive literature on its biochemical and genetic basis, we propose that color serves as an ideal trait for probing the evolutionary relationship between flower and fruit traits. To our knowledge, no previous study has used comparative methods to quantify the evolutionary dynamics (e.g., the number of transitions from one color to another) of flower and fruit colors across lineages, even though both traits have been studied in many individual clades (e.g., Armbruster 2002; S. D. Smith, Ané, and Baum 2008; Lomáscolo, Speranza, and Kimball 2008; Tripp et al. 2018; Lu et al. 2019; Sinnott-Armstrong et al. 2020; Hilgenhof et al. 2023).

Here, we present the first large-scale macroevolutionary analysis of flower and fruit color diversity to investigate broad patterns of variation across groups of plants as well as correlations in the tempo of color evolution. We grouped flower and fruit colors into eight categories each, to ensure comparability across the two organs. Using a dataset of 2989 species from 51 clades with both animal pollination and animal dispersal, and colors of flowers and fruits scored based on human vision for the same species, we estimate the evolutionary dynamics of color in terms of both transition rates and estimated transitions. As we detail below, our results reveal wide variation in color lability across organs and across clades, but also point to shared constraints on color evolution.

## METHODS

### DATA COLLECTION

#### Species/clade selection

We started with the list of species and fruit colors from Sinnott-Armstrong et al. (2018), which collated a dataset of human-described fruit colors spanning both the northern and southern hemispheres and all continents (except Antarctica). From that initial species list, we selected the subset of clades that are both animal-pollinated and animal-dispersed (Christenhusz, Fay, and Chase 2017). We supplemented those with three additional clades (*Gaultheria, Cornus,* and *Iochroma*), for which data was available from recent publications for fruit color (*Gaultheria, Cornus*) or flower color (*Iochroma*) for a majority of the species (see below). This procedure resulted in 51 clades distributed throughout the angiosperm phylogeny.

#### Phylogenies

For each of the 51 clades, we pruned a family-level phylogeny for that clade from “GBOTB.extended” tree (S. A. Smith and Brown 2018)). We visually examined the family-level phylogenies for each clade and removed 11 species that did not form a clade with the target taxon (e.g., two species of *Rhus* that group with other Anacardiaceae). We included an additional 175 species (and added corresponding color data, see below) to maximize the sampling within each clade. Because taxonomic names change frequently, we synonymized all names from our species list prior to extracting the trees using taxize() (Chamberlain et al. 2020).

#### Color data

The majority of fruit color data was derived from Sinnott-Armstrong et al. (2018), except for *Gaultheria* and *Cornus* which came from published datasets (Lu et al. 2019; Lindelof et al. 2020), and for *Iochroma* which came from field observations (S. D. Smith, pers. obs.). In addition to the species represented in Sinnott-Armstrong et al. (2018), we sought to obtain flower and fruit colors for all additional species represented in the pruned phylogenies. Specifically, we searched GBIF for each species and obtained either herbarium label descriptions (∼42% of species), research-grade iNaturalist images which we visually scored for color category from multiple images wherever possible (∼45% of species), and occasionally other sources (e.g., published studies, monographs, etc.). When images of living plants were available, those images were matched to a custom color chart to estimate the dominant color of the petals (see Fig. S1 for color scoring chart). For *Solanum,* we used data from Hilgenhof et al. (2023). In cases where flowers produced multiple colors, we used the primary display color as the color category for that species. In cases where fruits displayed multiple colors (e.g., between immature and mature fruits, or secondary structures) we used the color of the diaspore as the primary color.

In some cases, we were not able to find flower images or descriptions, and we removed such species from our dataset. In our final dataset, we included, on average, 29% of the species/clade, based on estimates of clade diversity from *Plants of the World* (Christenhusz et al 2017). To assess potential impacts of low sampling fractions for some clades on our results, we also ran our main analyses on subsets of clades which included more than 10% (43 clades), 20% (32 clades), 30% (18 clades), 40% (11 clades), or 50% (9 clades; see below for more details). In total, this procedure yielded a dataset with flower and fruit color along with inclusion in a phylogeny for 2989 species from 51 clades. Data will be made publicly available on Data Dryad (### provided upon acceptance).

#### Color categorization

In order to directly compare the number and rate of transitions in color between flowers and fruits, we categorized both fruit and flower color into eight distinct categories, in accordance with previous studies (Janson 1983; Fischer and Chapman 1993; Lu et al. 2019; Lindelof et al. 2020; Delmas, Kooyman, and Rossetto 2020; Wang et al. 2022 see Tables S1 and S2 for the number of species/clade in each color category). Although we used the same number of categories (eight for flowers, eight for fruits), we used different color categories between the two organs as some colors that are common in one organ are absent or nearly so in the other (e.g., fruits are often brown while flowers are rarely so). For fruits, we scored taxa according to eight categories commonly used by previous researchers (Janson 1983; Wheelwright and Janson 1985; Dominy, Svenning, and Li 2003; Duan, Goodale, and Quan 2014), which capture the most common human-visible colors: black, blue/purple, brown, green, yellow, orange, red, and white. The least common colors were grouped into one of these bins (e.g., blue also includes the less common purple and red includes the similarly rare pink). For flowers, we also adapted previously used schemes (e.g., Wang et al 2022, Dyer et al 2021, Delmas et al. 2020). Delmas et al. (2020) noted ten dominant colors, which we collapsed to eight categories for better comparison with fruit: blue/purple, dark (including black and other dark shades), green, orange, pink, red, white, and yellow) (see Supplemental Fig. S1 for more information on these color bins). Thus, the two organs share most categories (black/dark, blue/purple, green, yellow, orange, red, and white) except that flowers have a separate pink category (rare in fruits but common and ecologically important in flowers; Schemske and Bradshaw 1999) while fruits have a separate brown category (rare in flowers but common in some fruit clades and associated with mammal dispersal; Janson, 1983).

We did not attempt to account for UV reflectance in our data. UV reflectance is widespread in both flowers (Guldberg and Atsatt 1975; Koski 2020; Narbona et al. 2024) and fruits (Middleton et al. 2024; Willson and Whelan 1989), but is not visible to humans. Human vision captures the majority of color variation (Bergeron and Fuller 2018; Valenta et al. 2021) although cannot evaluate UV reflectance.

### STATISTICAL ANALYSES

#### Color diversity across clades

To identify whether clades tend to exhibit more color categories among their fruits or their flowers, we used a phylogenetic t-test (from the R package *phytools* (Revell 2024) to compare the number of categories across clades in the two organs. Then, to capture general patterns in the diversity of flower and fruit colors among our 51 clades, we calculated the Shannon diversity index for each clade based on the number of species per color category using the function diversity() from R package *vegan* (Oksanen et al. 2013). A high Shannon diversity indicates an even representation of different color categories in a clade, while a low Shannon diversity indicates an uneven representation (i.e., dominance of a single color).

#### Testing for correlation between number of transitions in flower and fruit colors

Due to the large number of categories (eight for each), we used stochastic character mapping with the “all rates different” model to estimate the number of transitions between color categories for flowers and fruits within each clade in the R package *phytools*. Because larger clades have more transitions in both flower and fruit color overall (Fig. S2), our null hypothesis was that the number of transitions would be equal between flowers and fruits (i.e., would be positively correlated along a 1:1 line). Thus, to test whether our data differed from that null distribution, we calculated the deviation from the 1:1 line, that is deviation from equal numbers of transitions as (# of fruit transitions – # of flower transitions). To account for differences in clade size, we scaled that deviation by the number of species in the clade as estimated by *Plants of the World (Christenhusz, Fay, and Chase 2017)*. In this model, a value for the deviation of 0.5 means that there is a gap between the number of transitions in fruits and flowers that is equal to 50% of the species in the clade, which represents a strong bias towards fruits. A deviation of −0.5 means that there are more flower transitions than fruit transitions, with the gap between those two numbers equaling 50% of the species in the clade. A deviation of 0 means that the same number of transitions was reconstructed in both flower and fruit color. Using these deviations, we then constructed a linear model with deviation as the response variable and the number of flower color transitions as the predictor variable in order to test for a relationship between flower color transitions and fruit color transitions across clades.

To evaluate whether our results were robust to different approaches, we performed two additional analyses to account for differences in clade size. First, we used the same deviations from the 1:1 line as above, but instead of scaling by clade size we included an interaction between the number of flower color transitions and the size of the clade. Second, we randomly subsampled 7 species from each of the 46 clades with greater than 7 species 100 times and generated stochastic character maps for each randomly subsampled phylogeny. Then, we counted the number of transitions in flower and fruit color on each of the subsampled phylogenies and averaged the number of transitions across all iterations. As above, we constructed a linear model with the deviation between fruit and flower color transitions as the response variable and the mean number of transitions in flowers as the predictor. We used a t-test to test whether the number of transitions in flower color and fruit color differed across clades, and an ANOVA to test for differences in the numbers of transitions in each organ between clades that had more flower color transitions (which we term “flower clades”) and those with more fruit color transitions (“fruit clades”).

#### Correlations between colors in flowers and fruits

To test whether any color category was positively correlated between flowers and fruits across clades (e.g., whether yellow fruits were more common in clades with more yellow flowers), we pruned the Smith and Brown (2018) phylogeny to one tip per clade to generate a clade-level phylogeny. We then calculated the proportion of species in each clade that produced each color category and used phylogenetic GLS in the R package caper (Orme, Freckleton, and Thomas, n.d.).

Next, we wanted to assess whether particular colors of flowers or fruits were associated with a clade being more biased towards flower or fruit transitions (e.g., if red — commonly associated with hummingbird pollination (Grant 1966; Bergamo et al. 2016; Altshuler 2003; Wessinger 2024) — were more likely to occur in lineages with elaborated flowers, e.g., flower clades). We used the clade-level phylogeny, combined with the proportion of species/clade producing each color. We then ran a phylogenetic GLS as described above to test whether any color categories were more common in “flower clades” or “fruit clades”.

#### Principal components analysis

We used principal components analysis on transition rates away from each color category in order to explore three aspects of color evolution: 1) whether particular colors are associated with being a “flower clade” or a “fruit clade” (e.g., are transition rates from white flowers high in flower clades but low in fruit clades?); 2) whether “flower clades” and “fruit clades” show similar correlations between color transitions (e.g., are transitions from red and blue-purple correlated in both organs?); and 3) whether there are correlations between flower color transition rates and fruit color transition rates across clades.

We estimated the rates of transition between color categories (8 categories total for both flowers and fruits) within each clade, for flowers and fruits separately, by extracting the estimated transition rate matrices from the stochastic character mapping. We then averaged the transition rates *from* each color category *to* all other categories in order to estimate the rate of transition from each color on average. We chose to use the estimated transition rates *from* each color category (rather than *to* each color category) because the originating color is the starting point for evolution and the transition rates directly predict persistence in any given state (Zanne et al. 2014). To counteract any effects of clade size on transition rates, we centered and scaled the transition rates by clade prior to input into the PC analysis.

We conducted three PCAs: one on fruit transitions alone, one on flower transitions alone, and one with both transition rates in the same analysis. We then used ANOVA to test whether “fruit clades” and “flower clades” cluster in different regions of PC space in each analysis.

## RESULTS

### Diversity of flower and fruit colors across clades

Most clades (34 of 51; 67%) exhibit more transitions in fruit color than in flower color (Fig. 1A). Fewer clades are biased towards flower color transitions (15 of 51; 29%) while two clades showed no difference in the number of the transitions in color in either organ. We divide clades into “fruit clades” (with more transitions in fruit color) and “flower clades” (with more transitions in flower color), and consequently ∼2/3 of clades are classified as “fruit clades” and ∼1/3 as “flower clades”. The mean number of color categories produced per clade was higher in fruits than in flowers (5.14 vs. 4.08, respectively; p = 0.004), and this difference was especially pronounced in fruit clades, which had more categories (mean: 5.54 categories/clade) compared with fruit colors in flower clades (3.89), flower colors in flower clades (4.53), or fruit colors in flower clades (4.20; p=0.009). According to the Shannon diversity index, *Diospyros* (8 colors across its 119 species) and *Capparis* (8 colors across 30 species) had the highest fruit color diversity, while *Ribes* had the highest flower color diversity (6 colors across 83 species; Fig. 1B,C; Supplemental Table S1). Five clades produced all eight fruit color categories (*Prunus, Capparis, Solanum, Passiflora,* and *Diospyros*) and eight clades produced seven of the fruit color categories (*Terminalia, Garcinia, Protium, Pouteria, Annona, Cissus,* and *Ribes*). In contrast, only two lineages produced all eight flower color categories (*Smilax* and *Pouteria*), and three lineages produced seven flower color categories (*Passiflora, Diospyros,* and *Cissus*).

**Figure 1.**
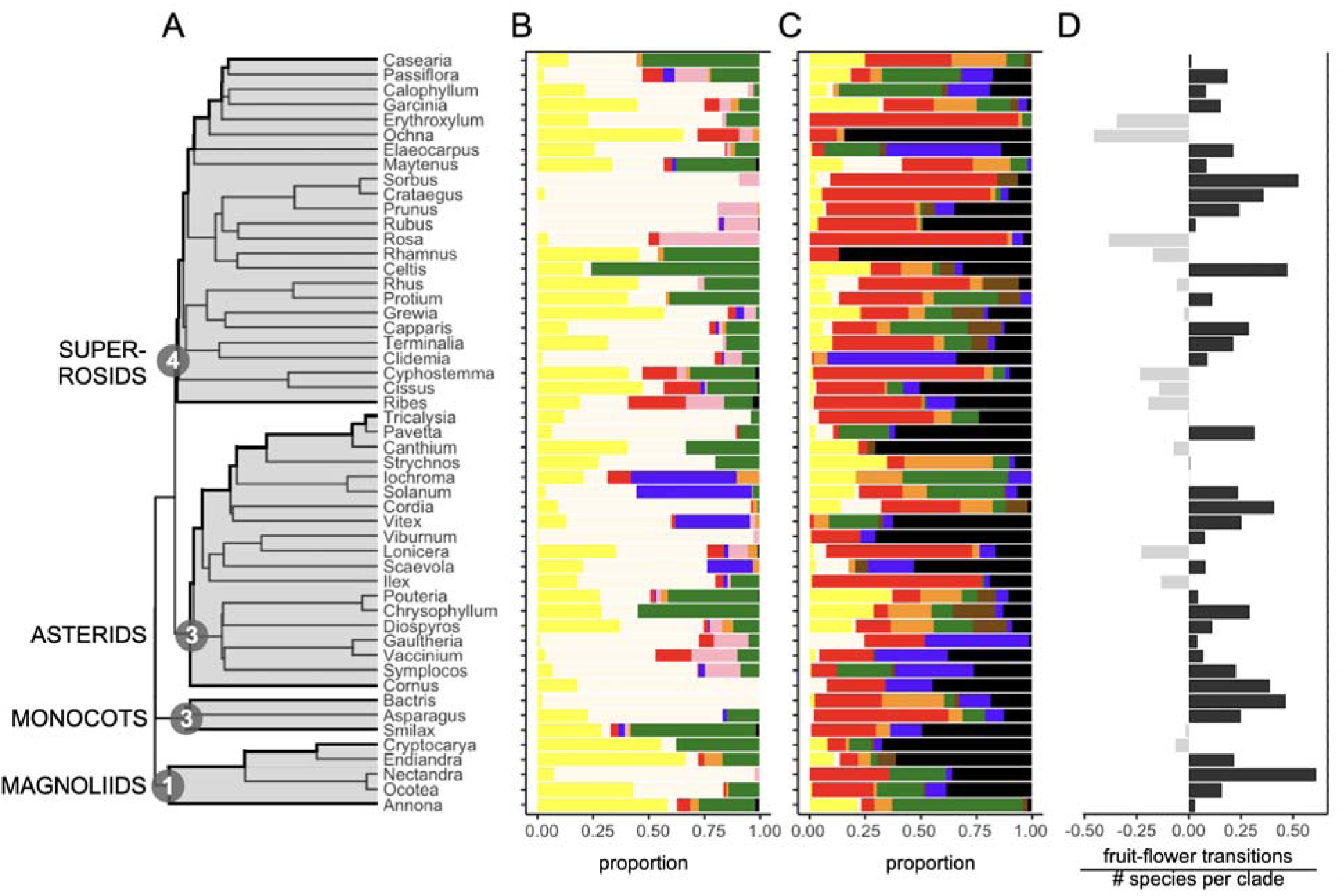
Patterns in flower and fruit color diversity across clades. A) Clades included in this study are distributed throughout the phylogeny of angiosperms, and include lineages from all of the major groups of plants (magnoliids, monocots, asterids, and superrosids). B) Proportion of species per clade in each flower color category (yellow, white [pale yellow in figure], red, blue-purple, pink, orange, green, black/dark). C) Proportion of species per clade in each fruit color category (yellow, white [pale yellow in figure], red, orange, green, brown, blue-purple, black/dark). D) The difference between the number of transitions in flower and fruit color, scaled by the number of species per clade in our dataset, reveals that most clades have more transitions in fruit color than in flower color. “Fruit clades” (difference > 0; black bars) have more transitions in fruit color, while “flower clades” (difference < 0; light gray bars) have more flower color transitions.

Overall, white was the most common color in flowers (48% of species) followed by yellow (16.9%), green (12.7%), blue/purple (9.9%), pink (7.0%), red (4.2%). Orange- (1.0%) and especially black/dark- (0.3%) flowered species were both exceedingly rare. In fruits, red was the most common color (30.2% of species), followed by black (21.3%). Green (13.2%), blue/purple (12.2%) and yellow (10.5%) were somewhat common, while orange (6.6%) and especially brown (3.0%) and white (2.9%) were rare.

### Negative correlation between flower and fruit color transitions across clades

We find a strong negative correlation in the number of color transitions that occur in flowers and fruits. Clades with many transitions in flower color have a correspondingly low number of transitions in fruit color, and vice versa (R^2^ = 0.48; p < 0.001; Fig. 2). This result is robust to sampling fraction such that the same negative correlation with a high R² (>0.46) is retained when filtering the dataset to only retain clades where we have at least 10% (43 clades), 20% (32 clades), 30% (18 clades), 40% (11 clades), or 50% (9 clades) of the total species included in our dataset (Fig. S3). When we include clade size as an interaction term with the number of flower transitions (rather than scaling transitions by the size of the clade), the interaction is highly significant (R^2^ = 0.75; p < 0.001). Our models using the subsampled phylogenies (Fig. S4) also found a strong correlation between the number of flower color transitions and the deviation in number of fruit color transitions (R^2^ = 0.40, p < 0.001).

**Figure 2.**
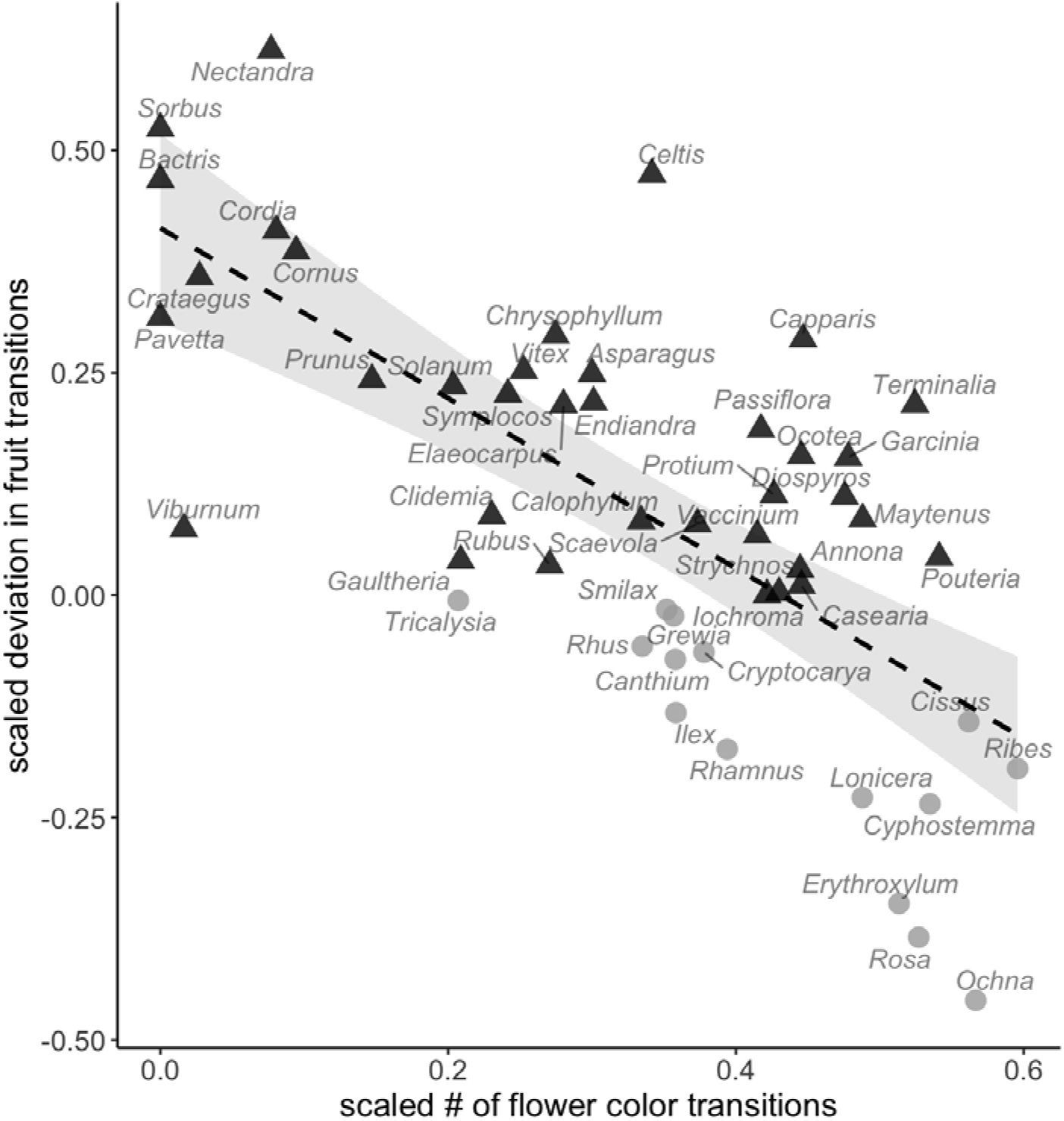
The number of transitions in flower color and fruit color are negatively correlated across clades. We compare the flower color transitions (scaled by the number of species per clade) against the scaled difference between fruit and flower transitions (scaled by the number of species per clade) to test whether the relationship between flower and fruit transitions differs from a slope of 1. There is a strong negative relationship (R^2^ = 0.48, p < 0.001) showing that some clades are strongly biased towards fruit color transitions (upper left) and other clades are strongly biased towards flower color transitions (lower right), which we name “fruit clades” (black triangles) and “flower clades” (gray circles), respectively.

#### Cross-organ correlations in color frequencies

The proportion of yellow-fruited species in any given clade was positively correlated with the proportion of yellow-flowered species (p=0.027, R^2^ = 0.10) in that clade, and similarly the proportion of orange-fruited species was positively correlated with the proportion of orange-flowered species (p=0.001, R^2^ = 0.20) when accounting for phylogenetic relatedness between clades. No other colors exhibited similar flower-fruit correlations (p > 0.05). The proportion of species/clade with red (p<0.001; R^2^ = 0.23) and yellow (p=0.002; R^2^=0.18) flowers were higher in “flower clades” with a greater bias toward flower color lability. Similarly, in fruits the proportion of species/clade with green (p=0.01; R^2^ = 0.11) and brown (p=0.03; R^2^=0.09) were more common in “fruit clades” with a greater bias towards fruit color lability.

### Ordination of transition rates

Flower and fruit color transition rates show similar correlations between colors in both organs. In flowers, PC1 and PC2 explain 49.3% and 15.2% of the variance, respectively, while in fruits PC1 and PC2 explain 54.1% and 16.1% of the variance, respectively. The color transition rates are positively correlated with PC1, suggesting that this axis represents overall magnitude of rates. The space occupied by flower and fruit clades overlaps for both organs (Fig. 3A,B), although flower transition rate space (as shown with the 90% ellipses in Fig. 3A) is larger for flower clades than for fruit clades. The converse is true for fruit transition rate space where the 90% ellipse for fruit clades is larger than for flower clades, (Fig. 3B). Nevertheless, both organs show some similar patterns of loading for the color categories. In particular, the transition rates from red, purple/blue and dark/black are closely aligned while other colors (except pink for flowers and green for fruits) load in the opposite direction on PC2 (Fig. 3). This pattern indicates that clades with high transition rates from purple/blue also have high rates for red and black/dark.

**Figure 3.**
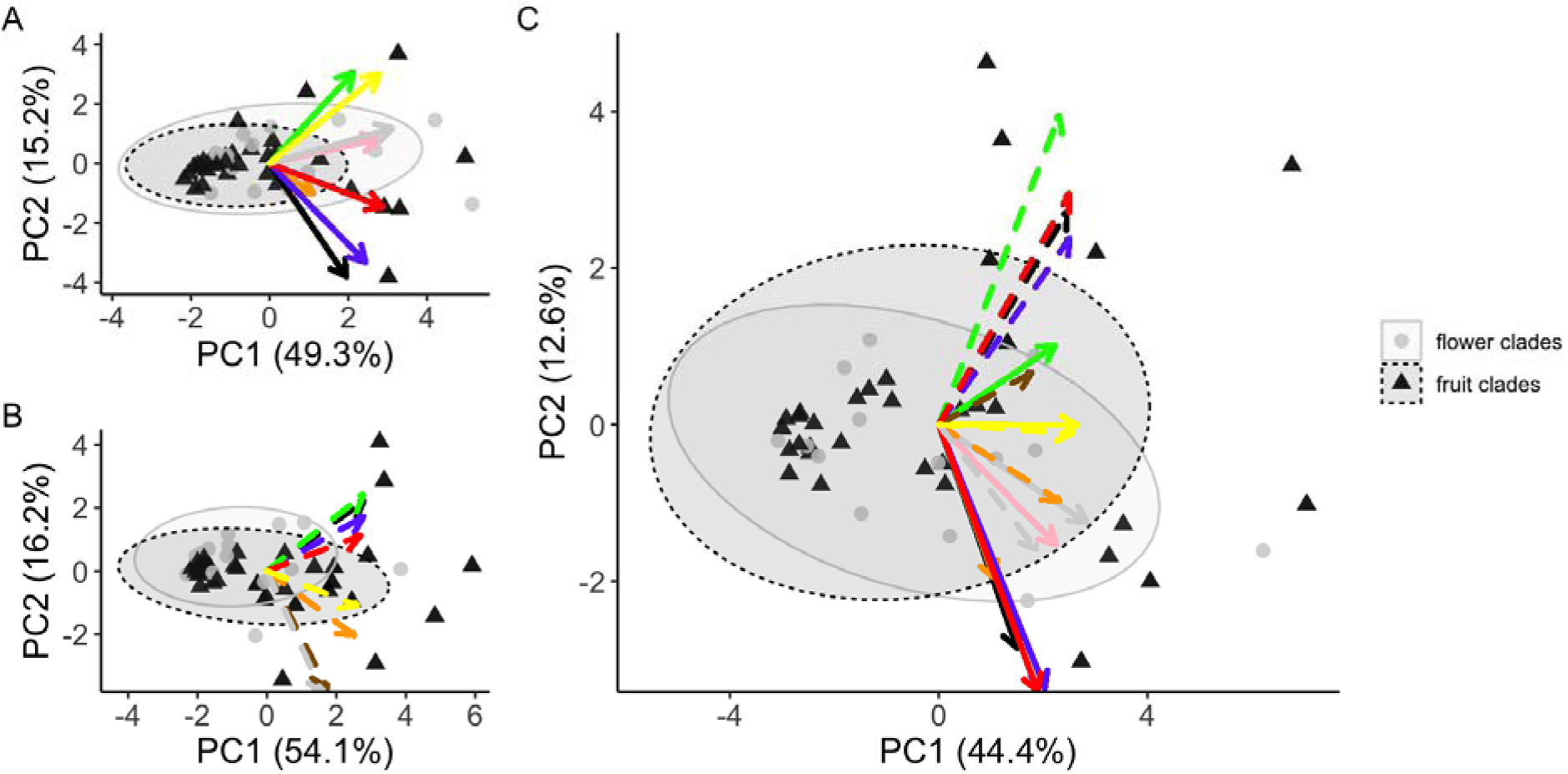
Principal components analyses of transition rates reveal consistent evolutionary patterns. In both (A) flowers and (B) fruits, loadings for transitions away from black/dark, red, and blue/purple are correlated within each organ. In neither organ is there any evidence that such patterns are driven by differences in “flower clades” or “fruit clades”. (C) When transition rates from both flowers and fruits are combined in the same analysis, transitions from black/dark, red, and blue/purple are again correlated within organs, but negatively correlated across organs (i.e., these colors are positively correlated with PC2 for fruits, but negatively correlated with PC2 for flowers). In contrast, yellow and to a lesser extent orange, green, and white are all positively correlated between organs, i.e., transitions from yellow in both flowers and fruits are positively correlated on PC1). Again, “flower clades” and “fruit clades” show no differences in location in PC space. Arrows correspond to the loading on each PC axis from a given color (e.g., red arrow = rate of transition away from red). Gray dots = “flower clades” and black triangles = “fruit clades”; 90% ellipses are drawn to show occurrence in PC space of flower clades (light gray) and fruit clades (dark gray).

When flower and fruit transition rates are included in the same analysis (Fig. 3c), we see that black/dark, red, and blue/purple again correlate within organ but load in opposite directions on PC2 between organs, such that clades with higher transition rates from black/dark, red, and blue/purple in flowers tend to have low transition rates away from those colors in fruits and vice versa. However, transition rates from yellow in both organs are positively correlated along PC1 and PC2, and similarly transition rates from green and white are positively correlated across the two organs. As with the individual organ analyses, flower and fruit clades are strongly overlapping (ANOVA, p > 0.1).

## DISCUSSION

Although flower color is often touted as one of the most diverse and labile traits in angiosperms (S. D. Smith and Goldberg 2015), our results reveal that, in most clades that rely on animals both for pollination and seed dispersal, fruit color is more labile than flower color. We also observed a macroevolutionary trade-off in the number of transitions in fruit colors and flower colors across clades, such that the two are negatively correlated. We term clades where flower color is highly labile but fruit color relatively static “flower clades”, while clades where fruit color evolves frequently but flower color exhibits evolutionary stasis “fruit clades”. Despite this contrast, we recover some connections between flower and fruit color diversity across clades, such as the tendency for yellow fruits to occur in clades with yellow flowers and the correlated transition rates for some color categories (e.g. red, purple/blue and black) in both organs. Below we discuss the possible underlying factors that may contribute to these intriguing macroevolutionary patterns.

### Flowers and fruits produce distinct distributions of colors

We find that flowers and fruits produce distinct distributions of colors, which are consistent with previous surveys (Delmas, Kooyman, and Rossetto 2020). The majority of flowers are white, yellow, or green, while black/dark and orange flowers are very rare (Delmas, Kooyman, and Rossetto 2020; Dyer et al. 2020). Most previous studies report ∼ 50% (or more, see Delmas, Kooyman, and Rossetto 2020) of species having white flowers (Dyer et al. 2020; Wang et al. 2022; Warren and Mackenzie 2001) which is in line with our finding of ∼48% of species having white flowers. Among fruits, black and red are by far the most common colors, constituting ∼50% of species in our dataset, again roughly in line with previous studies reporting 60-65% (Wheelwright and Janson 1985; Whitney 2009; Janson 1983; Sinnott-Armstrong et al. 2018). We also note that rare colors for each organ (especially red for flowers, and brown for fruit) tend to be more common in flower and fruit clades, respectively (Fig. 1).

We also observed that color distributions often appear clustered with respect to the phylogenetic relationships. For example, several closely related clades produce similar distributions of either fruit color (e.g., *Gaultheria, Symplocos, Vaccinium*; *Diospyros, Chrysophyllum, Pouteria*) or flower color (e.g., *Cyphostemma, Cissus, Ribes*). Rosaceae (here represented by *Sorbus, Prunus, Rosa, Rubus*, and *Crataegus*) have a somewhat unusual distribution of both fruit and flower colors: the majority of fruits are red (with a moderate frequency of black fruits, especially in *Prunus* and *Rubus*), and the flowers are almost exclusively white or pink. In addition, the majority of blue/purple flowers in our dataset fell within only four lineages, all of which are asterids and three of which (*Solanum, Iochroma,* and *Vitex*) are closely related (the fourth lineage is *Scaevola*). Such historical signals (i.e., ‘phylogenetic inertia’) are likely tied to the underlying biochemical pathways, which may facilitate some color transitions while constraining others (e.g., Wheeler et al. 2023).

While the distributions of color for each organ are similar across ours and other studies (e.g., white being the most common flower color), conclusions about the relative diversity of flower and fruit colors varies. Here we found that the number of color categories per clade was slightly (but significantly) higher in fruits than in flowers. Stournaras et al. (2013) report the opposite pattern, although they focused on bird-dispersed fruits, which largely excludes colors associated with mammal dispersal (yellow, orange, green, brown; Janson 1983; Fischer and Chapman 1993). Whitney (2009) also reported higher diversity of flowers than fruits, but incorporated physical traits (size) in addition to color to estimate the morphospace occupied by each organ. Both of these studies, like ours, focused on animal-pollinated and animal-dispersed fruits, and we expect additional differences in patterns of color diversity if analyses were expanded to abiotically-pollinated or dispersed clades. For example, our dataset included only 12 blue-flowered species (out of ∼3000; 0.4%), hence our lumping of blue with purple for analyses. The survey of Dyer et al. (2020) reports a much higher fraction (5-10% of species having blue flowers), but importantly they include dry-fruited lineages, suggesting that dry-fruited but animal-pollinated lineages may have different distributions of flower colors than the lineages that we include here that rely on animals for both pollination and seed dispersal.

One possible explanation for these differences in color distributions between flowers and fruits lies in the different animal guilds involved in pollination and seed dispersal. The visual systems of pollinators (i.e., bees, birds, bats, beetles, wasps) and dispersers (i.e., birds, mammals) may select for different color palettes. In fruits, blue, black and red fruits tend to be dispersed by birds while yellow, orange, green and brown are commonly associated with mammal dispersal, although such patterns are not universal (Janson 1983; Fischer and Chapman 1993; Valenta, Nevo, and Chapman 2018). Mammals are common dispersers especially in the tropics (Sinnott-Armstrong, Donoghue, and Jetz 2021; Losada, Suárez-Couselo, and Sobral 2024), but they are comparatively rare as pollinators (more than 2/3 of angiosperms are insect-pollinated (Rodger et al. 2021; Stephens et al. 2023). Moreover, unlike in flowers, fruit color-producing pigments can also act as a reward directly. The blue-purple and red anthocyanins commonly found in fruits have strong antioxidant activity and may be selected for by dispersers (Schaefer, Rentzsch, and Breuer 2008; Jiménez-Gallardo et al. 2025). Finally, flowers and fruits may differ in the structure of their interaction networks, with plant-pollinator networks often exhibiting specialization to ensure transfer between conspecifics (Chittka, Thomson, and Waser 1999; Gumbert, Kunze, and Chittka 1999; Wheelwright and Orians 1982) and plant-disperser networks being more generalized to move seeds more broadly and to a larger range of microhabitats (Wheelwright and Orians 1982; Harms et al. 2000; Russo and Augspurger 2004).

### Negative correlation between flower and fruit color lability

Our joint analysis of flower and fruit colors across clades revealed an unexpected trade-off in evolutionary lability, namely that clades with many transitions in fruit color tend to have fewer transitions in flower color, and vice versa. This pattern is exemplified by many well-known clades: for instance, *Viburnum* has four distinct fruit colors with at least 9 transitions (Sinnott-Armstrong et al. 2020) in color, while the vast majority of *Viburnum* species’ flowers are white with only a few that are pink (Fig. 1). Notably, both “flower clades” and “fruit clades” include a mixture of large and small clades, e.g., relatively large clades such as *Elaeocarpus* (fruit clade) and *Cissus* (flower clade), and relatively small clades such as *Cornus* (fruit clade) and *Iochroma* (flower clade). While the cause(s) of this negative correlation are uncertain, we have identified two hypotheses, the first related to resource allocation and the second to external environmental factors, which we discuss below.

#### Hypothesis 1: Differential resource allocation throughout plant reproduction

Plants are fundamentally limited in the resources they have available to allocate to the different stages of reproduction (Roddy et al. 2021), which may lead to differing strategies of investment of those resources into reproductive organs to ensure reproductive success (Charnov 1987; Lloyd 1987; Roff and Fairbairn 2007). Trade-offs in resource allocation to different functions have an extensive literature for vegetative traits (i.e., the leaf economics spectrum, Wright et al. 2004), and recently, an analogous economics spectrum has been proposed for flowers (Roddy et al. 2021). In reproductive traits, the best-studied relationships are trade-offs between size and number, which has been reported in flowers (Cohen and Dukas 1990; Kettle et al. 2011; Caruso, Maherali, and Benscoter 2012; Cao and Worley 2013; Vasconcelos and Proença 2015), ovules (Greenway and Harder 2007), and seeds (e.g., Jakobsson and Eriksson 2000; Bentos et al. 2014). Although such trade-offs between flowers and fruits have not been well-studied, theoretical work (e.g., Morgan 1993) suggests that such trade-offs should be expected. As one example, the “leaf intensity premium” hypothesis suggests that species that produce large leaves should produce fewer of them, and as a consequence fewer flowers and fruits due to limited opportunities to convert vegetative buds into reproductive shoots (Kleiman and Aarssen 2007; E-Vojtkó et al. 2022).

Differential investment in reproductive effort, in line with the “economics spectra” ideas, may explain the trade-off that we observe between flower and fruit color lability. Investment in flowers through evolving to better attract local pollinators, for example, may increase fertilization rate; if fertilization results in many fruits, selection on fruit traits may be reduced because there are plenty of fruits available to achieve reproductive success. On the other hand, if pollination rarely succeeds, the dispersal of individual fruits becomes critical for fitness, and selection towards diversifying fruit traits to better attract dispersers may occur. Such divergent investment strategies may explain the occurrence of “flower clades” and “fruit clades”. If differential investment into either flowers or fruits indeed is a common pattern across angiosperms, we may expect the most extreme cases of “flower clades” and “fruit clades” in abiotically dispersed and abiotically pollinated lineages, respectively. Anecdotally, such a pattern does indeed seem to occur among “flower clades”: many lineages with the most elaborated flowers (e.g., Orchidaceae*, Aquilegia, Mimulus, Penstemon,* some Melastomataceae, and others (Christenhusz et al., 2017)) have dry, wind-dispersed fruits.

The costs of pigment production may contribute to the trade-offs that we observed. In fruits, chlorophyll (and corresponding green coloration) in immature fruits can photosynthesize significant fractions of the carbon needed to maintain a fruit during its maturation (Bazzazz, Carlson, and Harper 1979), which can take many weeks to months. The off-setting of the cost of fruit production via *in situ* photosynthesis may facilitate the exploration and production of a wider range of pigments in mature fruits, where such pigments also perform other functions (e.g., defense against pathogens; Schaefer, Rentzsch, and Breuer 2008). In flowers, in contrast, chlorophyll in petals is unlikely to contribute substantially to the costs of flower production, because petals usually remain in bud for most of their development and, in the vast majority of species, are displayed for <10 days (Ashman and Schoen 1996). Consequently, white may represent a low investment with either a lack of pigments or a restricted set of UV-absorbing compounds (e.g. flavones, flavonols; Kazuma, Noda, and Suzuki 2003; Narbona et al. 2024). Green petals are common in flowers, as we confirm here, and may represent a compromise between contribution to photosynthetic capacity and minimizing costs (e.g., reduced signaling effectiveness due to lower contrast with background foliage; Del Valle et al. 2024). If so, we would expect that longer-lived petals would be more likely to exhibit green color due to chlorophyll contributing to the carbon costs of petal maintenance (as long-lived flowers can incur a substantial water- and energetic cost (Nobel 1977; Ashman and Schoen 1996). Overall, understanding the possible role of energetic investments in shaping flower and fruit color variation (and co-variation) will require detailed studies of the costs and benefits of pigment production (Chalker-Scott 1999).

#### Hypothesis 2: Habitat type and the diversity of pollinators and dispersers

Extrinsic biotic factors could select for opposing patterns of color diversity between the two phases of reproduction, leading to the observed negative correlation. One possible factor is habitat type (Givnish et al. 2005, 2020): pollinator visitation rates and diversity can be higher in open habitats and canopy gaps (Herrera 1995; Walters and Stiles 1996; Eckerter et al. 2019; McCabe et al. 2019; Mullally et al. 2019), perhaps due to warmer temperatures encouraging more insect (particularly bee) pollinators (Orr et al. 2021). The availability of a diverse suite of pollinators is likely a prerequisite for high color lability in “flower clades”, especially as shifts in pollinators are well-known to be associated with shifts in floral colors (S. D. Smith, Ané, and Baum 2008; Tripp and Manos 2008; Moré et al. 2020). Similarly, disperser diversity may be a prerequisite to fruit color lability in “fruit clades”. Fleshy fruits are more common in species occurring under closed canopies (Givnish et al. 2005), and both fleshy fruits and frugivores are often more abundant at forest edges (Menke, Böhning-Gaese, and Schleuning 2012) and in higher canopy strata (Schleuning et al. 2011; Thiel et al. 2023). Biome and habitat type are fairly conserved in most plant lineages (Edwards and Donoghue 2013; Donoghue and Edwards 2014), which may explain why some lineages may have labile flower colors (due to greater pollinator diversity in the biome in which that lineage diversified) while other lineages tend to favor diversification of fruit color (when such lineages primarily occupy biomes with high disperser diversity). However, at this point, the relationship between habitat type and color lability remains speculative.

### Color evolution along lines of least resistance

Our finding that clades that experience transitions involving blue/purple also tend to experience transitions from red and black (Fig. 3) points to an underlying role for pigment pathways in shaping the color diversity across clades and organs. These hues are typically produced in flowers and fruit by the blue, purple and red anthocyanin pigments (e.g., black/dark, red, blue/purple Chenery 1948; Grotewold 2006; Schaefer, Rentzsch, and Breuer 2008; e.g., black/dark, red, blue/purple; Tanaka, Sasaki, and Ohmiya 2008; Ranganath 2022); although other compounds (e.g., carotenoids, betalains) as well as structural colors can also be involved (e.g., Ng and Smith 2016; Sinnott-Armstrong et al. 2023). Thus, shifts between blue/purple and red can be achieved simply through an alteration of the type of anthocyanin produced, offering an appealing explanation for the positive correlated transition rates observed here (Fig. 3).

Interestingly, clades tend to either use these colors in their flowers or in their fruits, but less often do they use the same colors in both organs (i.e., blue/purple flowers in *Solanum, Iochroma, Vitex, Scaevola*, but blue/purple fruits in *Clidemia, Gaultheria, Symplocos, Vaccinium*; Fig. 3C).

In addition, the combined PC analysis indicated correlations of other color categories across organs. For example, yellow transition rates are tightly aligned between flowers and fruits, and green and orange load in similar manners on both PC axes (Fig. 3C). These three colors commonly derive from chlorophylls and carotenoids that accumulate in hydrophobic plastids (chloroplasts or chromoplasts, respectively), and the similarity of their evolutionary dynamics across clades and organs is consistent with their deep evolutionary history. Congruent with this evolutionary pattern, Dellinger et al. (2025) found that both yellow-flowered and yellow-fruited species were more common in arid habitats, suggesting a possible physiological explanation. For instance, in addition to their role in color, carotenoids also participate in desiccation resistance in red algae (Zhao et al. 2022), cyanobacteria (Yang et al. 2019), and vascular plants (Augusti et al. 2001; Vieira et al. 2024). Overall, the physiological importance of chlorophylls and carotenoids may help to explain the wide distribution of green and yellow flowers as well as green, yellow and orange fruits across clades (Fig. 1) and in turn, their similar patterns of color transitions (Fig. 3C).

### Limitations and future directions

Although we endeavored to broadly sample plant lineages, there are several limitations that — if overcome — would enhance our understanding of the relationship between flower and fruit evolution. First, color categories, as perceived by humans, capture a substantial portion of the variation in colors in nature (Bergeron and Fuller 2018; Valenta et al. 2021) but do not fully capture color as perceived by the relevant animals (pollinators and dispersers) and omit any UV reflectance. UV reflection can contribute significantly to the color signal in both flowers and fruits albeit through different mechanisms (Koski 2020; Middleton et al. 2024). In addition, our color classification scheme did not capture multi-colored displays, which are important sources of color contrast (i.e., stamens in flowers, pedicels or immature fruits in fruits (Wheelwright and Janson 1985; de Camargo et al. 2015). Inclusion of these additional aspects of signal diversity would provide a more complete understanding of the relationship between flower and fruit color evolution as well as the relationship between color and animal mutualists.

## CONCLUSIONS

While previous comparative studies of flower and fruit colors have focused on patterns of diversity (e.g., Whitney et al. 2009; Stournaras et al. 2013), we took an explicitly phylogenetic approach to investigating the dynamics of color evolution in both flowers and fruits, within the same species and clades. The presence of diverse colors within a clade is only part of the picture of color evolution; color lability (the frequency of transitions) is also a major factor. Our results indicate that fruit colors are only slightly more diverse than flower colors in any given clade, however the number of transitions in fruit colors is much higher on average and concentrated in particular clades (‘fruit clades’). The negative correlation between flower and fruit color transitions across clades suggest that “flower clades” and “fruit clades” pursue different strategies to achieving reproductive success. Despite this apparent trade-off in color evolution between flowers and fruits, both organs rely on the same biochemical components for producing color, which may explain correlated trends in transition rates for classes of colors that typically derive from the same classes of pigments. Thus, while plant organs have been historically treated as distinct modules that evolve relatively independently, our findings suggest that flowers and fruits may not be as modular as previously thought. Environmental factors, investment strategies, and cross-organ developmental constraints may all interact to generate the correlations described here, and better understanding these factors will be critical for explaining large-scale, macroevolutionary patterns in angiosperm diversity.

## ACKNOWLEDGEMENTS

We gratefully acknowledge helpful discussions with Thais Vasconcelos and Michael Donoghue. Funding for MSA and LM came from NSF DBI-1907293. Funding for ASD came from Austrian Science Fund FWF T-1186, and for SS from NSF-DEB 1553114. We are also immensely grateful to Justen Whittall and two anonymous reviewers, whose comments greatly improved the manuscript.

