## Supplemental Material for "Flower clades and fruit clades: trade-offs in color diversification across angiosperms"

**Supplemental Figure S1.** Color scoring chart used to characterize the colors of flowers from images from iNaturalist, GBIF, and occasionally other sources. We grouped colors into eight categories for flowers: white (w1), black/dark (all of the darkest categories, i.e., w5, b5, g5, y5, o5, r5, pu5, pi5, t5), pink (pi1, pi2, pi3, pi4), purple/blue (pu1, pu2, pu3, pu4, b1, b2, b3, b4), yellow (y1, y2, y3, y4), orange (o1, o2, o3, o4), red (r1, r2, r3, r4), and green (g1, g2, g3, g4).

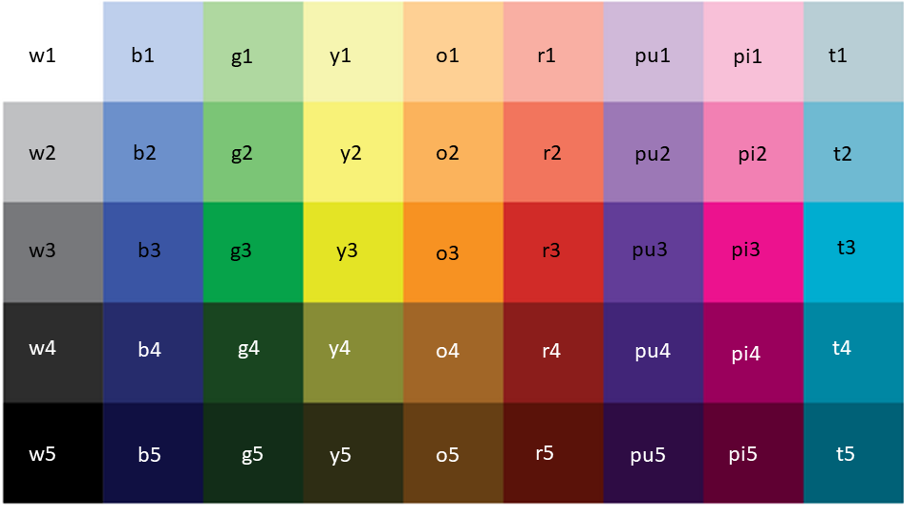

**Supplemental Figure S2:** Larger clades have a larger number of transitions in both (A) flower and (B) fruit colors. Counts of transitions are derived from the mean number of transitions from 100 stochastic maps for each clade, while the number of species in the clade is estimated from *Plants of the World* (as described in the main text).

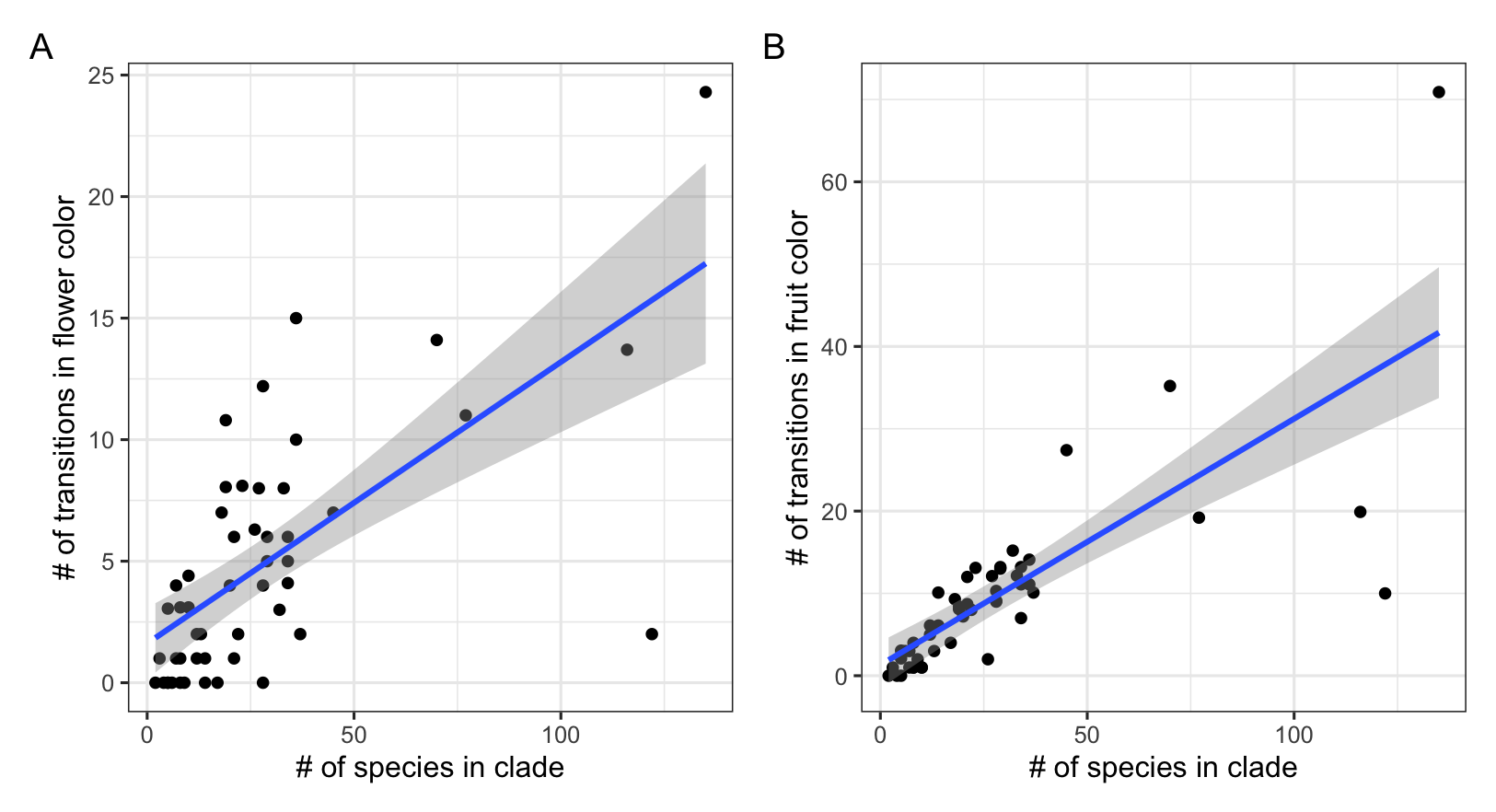

**Supplemental Figure S3**. To assess whether the degree of species coverage (i.e., the number of species/clade in our dataset and the estimated total richness of that clade, according to *Plants of the World*) affected our results, we tested whether there was a negative correlation between flower and fruit transitions if we filtered to only include clades with A) all clades [same species included as in the main text], B) 10% coverage, C) 20% coverage, D) 30% coverage, E) 40% coverage, and F) 50% coverage. In all cases, there was a negative correlation which was statistically significant with a high R^2^ (> 0.48).

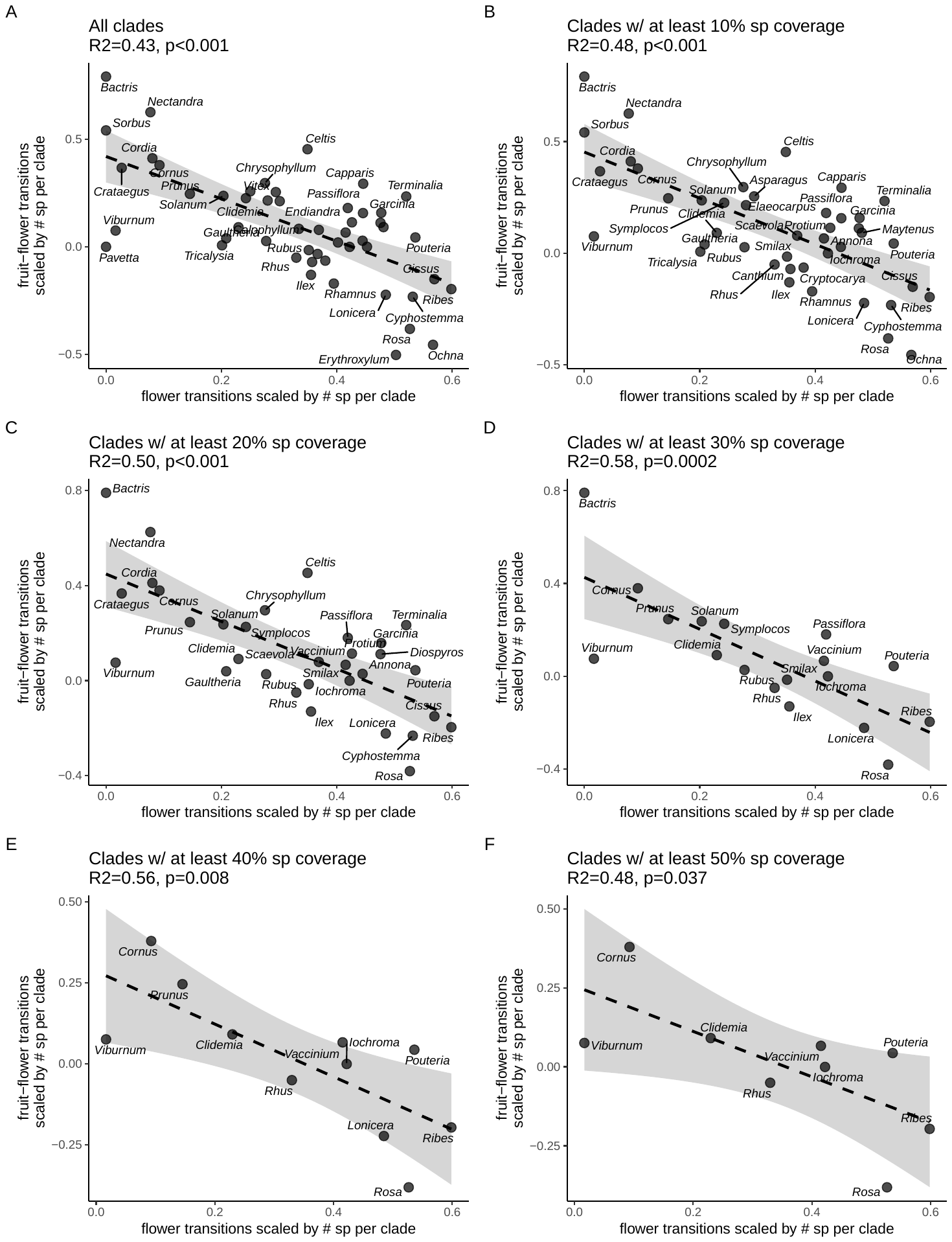

**Supplemental Figure S4.** Negative correlation between flower and fruit color transitions across clades based on a subsampling approach. As an alternative to the main analysis in the text, we subsampled each phylogeny down to 7 species 100 times, then used stochastic character mapping to estimate the total number of transitions in color in both flowers and fruits. We find a negative correlation (R^2^ = 0.40, p < 0.001) such that clades with more flower color transitions tend to have fewer fruit color transitions.

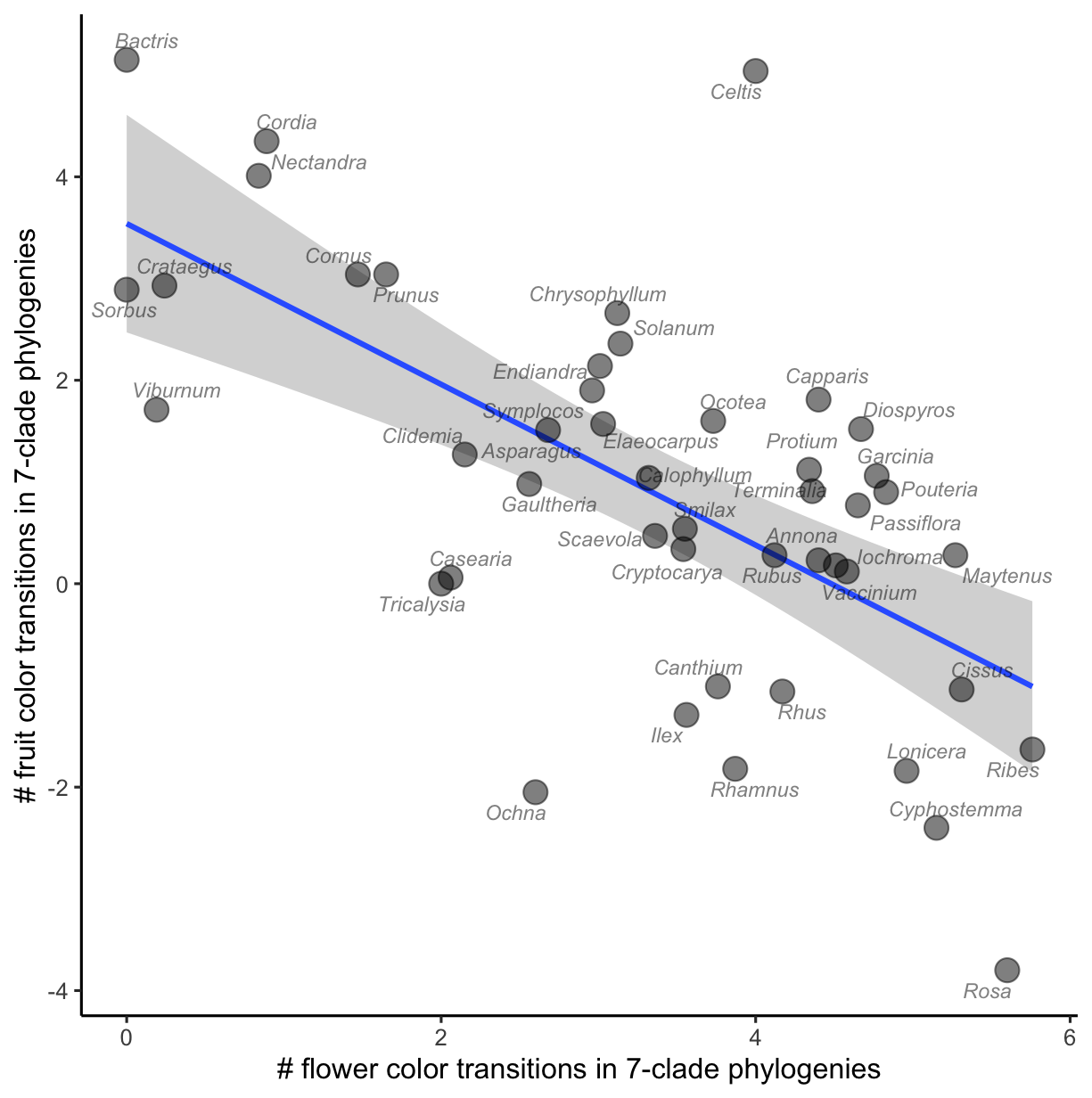

**Supplemental Table S1.** The number of species in each color category, per clade, for fruits, after classification and grouping into eight color categories (black, blue, brown, green, orange, red, white, yellow).

| **clade** | **black** | **blue** | **brown** | **green** | **orange** | **red** | **white** | **yellow** | **total** |
| --- | --- | --- | --- | --- | --- | --- | --- | --- | --- |
| *Annona* | 1 | 0 | 1 | 19 | 4 | 3 | 1 | 8 | **37** |
| *Asparagus* | 6 | 4 | 0 | 2 | 2 | 18 | 1 | 0 | **33** |
| *Bactris* | 5 | 4 | 1 | 0 | 5 | 12 | 0 | 1 | **28** |
| *Calophyllum* | 2 | 0 | 1 | 6 | 0 | 0 | 0 | 3 | **12** |
| *Canthium* | 10 | 0 | 1 | 0 | 0 | 0 | 0 | 3 | **14** |
| *Capparis* | 2 | 1 | 8 | 9 | 4 | 7 | 2 | 2 | **35** |
| *Casearia* | 0 | 0 | 1 | 0 | 4 | 4 | 0 | 0 | **9** |
| *Celtis* | 1 | 0 | 2 | 1 | 3 | 2 | 0 | 3 | **12** |
| *Chrysophyllum* | 4 | 0 | 5 | 1 | 3 | 2 | 0 | 6 | **21** |
| *Cissus* | 37 | 5 | 0 | 4 | 1 | 24 | 1 | 2 | **74** |
| *Clidemia* | 27 | 51 | 0 | 0 | 4 | 0 | 1 | 0 | **83** |
| *Cordia* | 1 | 0 | 7 | 5 | 9 | 28 | 15 | 11 | **76** |
| *Cornus* | 16 | 8 | 0 | 0 | 0 | 10 | 3 | 0 | **37** |
| *Crataegus* | 7 | 3 | 0 | 1 | 0 | 27 | 0 | 1 | **39** |
| *Cryptocarya* | 14 | 2 | 0 | 6 | 0 | 1 | 0 | 4 | **27** |
| *Cyphostemma* | 3 | 1 | 0 | 3 | 2 | 22 | 0 | 0 | **31** |
| *Diospyros* | 10 | 2 | 16 | 20 | 27 | 20 | 2 | 22 | **119** |
| *Elaeocarpus* | 11 | 32 | 3 | 15 | 0 | 5 | 0 | 0 | **66** |
| *Endiandra* | 4 | 0 | 2 | 1 | 0 | 2 | 0 | 1 | **10** |
| *Erythroxylum* | 0 | 0 | 0 | 0 | 1 | 5 | 0 | 0 | **6** |
| *Garcinia* | 0 | 3 | 2 | 11 | 12 | 11 | 1 | 20 | **60** |
| *Gaultheria* | 1 | 36 | 0 | 0 | 0 | 21 | 19 | 0 | **77** |
| *Grewia* | 1 | 0 | 0 | 0 | 0 | 2 | 0 | 0 | **3** |
| *Ilex* | 24 | 4 | 0 | 1 | 0 | 99 | 0 | 2 | **130** |
| *Iochroma* | 0 | 2 | 0 | 9 | 4 | 0 | 0 | 4 | **19** |
| *Lonicera* | 14 | 6 | 0 | 0 | 2 | 56 | 4 | 1 | **83** |
| *Maytenus* | 0 | 1 | 0 | 2 | 4 | 14 | 7 | 7 | **35** |
| *Nectandra* | 9 | 1 | 0 | 9 | 0 | 7 | 0 | 0 | **26** |
| *Ochna* | 8 | 0 | 0 | 0 | 0 | 1 | 0 | 0 | **9** |
| *Ocotea* | 15 | 8 | 0 | 12 | 1 | 8 | 0 | 1 | **45** |
| *Passiflora* | 36 | 28 | 2 | 60 | 9 | 14 | 1 | 36 | **186** |
| *Pavetta* | 9 | 2 | 0 | 1 | 0 | 1 | 0 | 0 | **13** |
| *Pouteria* | 11 | 8 | 8 | 8 | 23 | 17 | 0 | 44 | **119** |
| *Protium* | 0 | 3 | 6 | 11 | 3 | 18 | 1 | 5 | **47** |
| *Prunus* | 30 | 9 | 5 | 1 | 3 | 37 | 1 | 7 | **93** |
| *Rhamnus* | 20 | 0 | 0 | 0 | 0 | 5 | 0 | 0 | **25** |
| *Rhus* | 0 | 0 | 1 | 0 | 3 | 24 | 7 | 2 | **37** |
| *Ribes* | 27 | 12 | 0 | 1 | 1 | 40 | 1 | 1 | **83** |
| *Rosa* | 4 | 5 | 0 | 0 | 2 | 76 | 0 | 0 | **87** |
| *Rubus* | 34 | 0 | 0 | 0 | 2 | 49 | 0 | 5 | **90** |
| *Scaevola* | 18 | 7 | 2 | 0 | 1 | 0 | 5 | 1 | **34** |
| *Smilax* | 44 | 15 | 0 | 0 | 3 | 29 | 1 | 0 | **92** |
| *Solanum* | 24 | 22 | 5 | 139 | 49 | 92 | 9 | 102 | **442** |
| *Sorbus* | 0 | 0 | 2 | 0 | 0 | 10 | 2 | 0 | **14** |
| *Strychnos* | 1 | 0 | 0 | 0 | 3 | 0 | 0 | 3 | **7** |
| *Symplocos* | 18 | 37 | 1 | 28 | 0 | 11 | 0 | 1 | **96** |
| *Terminalia* | 5 | 1 | 5 | 6 | 2 | 18 | 0 | 4 | **41** |
| *Tricalysia* | 4 | 0 | 0 | 1 | 0 | 5 | 0 | 0 | **10** |
| *Vaccinium* | 31 | 31 | 1 | 0 | 0 | 20 | 2 | 3 | **88** |
| *Viburnum* | 88 | 10 | 0 | 0 | 0 | 26 | 0 | 1 | **125** |
| *Vitex* | 2 | 0 | 0 | 1 | 1 | 0 | 0 | 0 | **4** |
| ***TOTAL*** | **639** | **364** | **88** | **394** | **197** | **903** | **87** | **317** | **2989** |

**Supplemental Table S2.** The number of species in each color category, per clade, for flowers, after classification and grouping into eight color categories (black/dark, green, orange, pink, purple, red, white, yellow).

| **clade** | **black** | **green** | **orange** | **pink** | **purple** | **red** | **white** | **yellow** | **total** |
| --- | --- | --- | --- | --- | --- | --- | --- | --- | --- |
| *Annona* | 1 | 9 | 2 | 0 | 0 | 3 | 1 | 21 | **37** |
| *Asparagus* | 0 | 4 | 0 | 0 | 0 | 0 | 21 | 8 | **33** |
| *Bactris* | 0 | 0 | 0 | 0 | 0 | 0 | 28 | 0 | **28** |
| *Calophyllum* | 0 | 1 | 0 | 0 | 0 | 0 | 8 | 3 | **12** |
| *Canthium* | 0 | 3 | 0 | 0 | 0 | 0 | 4 | 7 | **14** |
| *Capparis* | 0 | 9 | 1 | 0 | 0 | 0 | 20 | 5 | **35** |
| *Casearia* | 0 | 5 | 1 | 0 | 0 | 0 | 3 | 0 | **9** |
| *Celtis* | 0 | 8 | 0 | 0 | 0 | 0 | 0 | 4 | **12** |
| *Chrysophyllum* | 0 | 12 | 0 | 0 | 0 | 0 | 2 | 7 | **21** |
| *Cissus* | 1 | 16 | 0 | 1 | 2 | 13 | 9 | 32 | **74** |
| *Clidemia* | 0 | 7 | 0 | 7 | 1 | 2 | 64 | 2 | **83** |
| *Cordia* | 0 | 1 | 0 | 0 | 0 | 1 | 67 | 7 | **76** |
| *Cornus* | 0 | 0 | 0 | 0 | 0 | 0 | 30 | 7 | **37** |
| *Crataegus* | 0 | 0 | 0 | 0 | 0 | 0 | 38 | 1 | **39** |
| *Cryptocarya* | 0 | 11 | 0 | 0 | 0 | 0 | 1 | 15 | **27** |
| *Cyphostemma* | 0 | 8 | 0 | 2 | 0 | 7 | 2 | 12 | **31** |
| *Diospyros* | 0 | 12 | 6 | 8 | 1 | 3 | 43 | 46 | **119** |
| *Elaeocarpus* | 0 | 6 | 1 | 2 | 0 | 0 | 45 | 12 | **66** |
| *Endiandra* | 0 | 1 | 1 | 0 | 0 | 0 | 1 | 7 | **10** |
| *Erythroxylum* | 0 | 2 | 0 | 1 | 0 | 0 | 3 | 0 | **6** |
| *Garcinia* | 0 | 5 | 3 | 3 | 0 | 5 | 19 | 25 | **60** |
| *Gaultheria* | 0 | 4 | 0 | 12 | 0 | 5 | 55 | 1 | **77** |
| *Grewia* | 0 | 0 | 0 | 0 | 1 | 0 | 2 | 0 | **3** |
| *Ilex* | 0 | 15 | 0 | 2 | 3 | 4 | 83 | 23 | **130** |
| *Iochroma* | 0 | 0 | 2 | 0 | 9 | 2 | 2 | 4 | **19** |
| *Lonicera* | 0 | 0 | 4 | 7 | 2 | 6 | 34 | 30 | **83** |
| *Maytenus* | 1 | 12 | 0 | 0 | 1 | 2 | 10 | 9 | **35** |
| *Nectandra* | 0 | 0 | 0 | 0 | 0 | 0 | 24 | 2 | **26** |
| *Ochna* | 0 | 0 | 0 | 1 | 0 | 2 | 2 | 4 | **9** |
| *Ocotea* | 0 | 4 | 0 | 0 | 0 | 0 | 20 | 21 | **45** |
| *Passiflora* | 0 | 43 | 2 | 29 | 7 | 17 | 83 | 5 | **186** |
| *Pavetta* | 0 | 0 | 0 | 0 | 0 | 0 | 13 | 0 | **13** |
| *Pouteria* | 1 | 48 | 4 | 3 | 1 | 3 | 29 | 30 | **119** |
| *Protium* | 0 | 18 | 0 | 0 | 0 | 0 | 8 | 21 | **47** |
| *Prunus* | 0 | 0 | 0 | 17 | 0 | 0 | 76 | 0 | **93** |
| *Rhamnus* | 0 | 6 | 1 | 0 | 0 | 0 | 3 | 15 | **25** |
| *Rhus* | 0 | 7 | 0 | 2 | 0 | 0 | 13 | 15 | **37** |
| *Ribes* | 3 | 11 | 0 | 13 | 0 | 23 | 17 | 16 | **83** |
| *Rosa* | 0 | 0 | 0 | 37 | 0 | 5 | 40 | 5 | **87** |
| *Rubus* | 1 | 0 | 0 | 19 | 3 | 1 | 66 | 0 | **90** |
| *Scaevola* | 0 | 0 | 1 | 0 | 7 | 0 | 19 | 7 | **34** |
| *Smilax* | 2 | 50 | 1 | 2 | 2 | 3 | 3 | 29 | **92** |
| *Solanum* | 0 | 12 | 0 | 2 | 251 | 0 | 158 | 19 | **442** |
| *Sorbus* | 0 | 0 | 0 | 0 | 0 | 0 | 14 | 0 | **14** |
| *Strychnos* | 0 | 2 | 0 | 0 | 0 | 0 | 4 | 1 | **7** |
| *Symplocos* | 0 | 9 | 0 | 17 | 1 | 1 | 62 | 6 | **96** |
| *Terminalia* | 0 | 8 | 0 | 1 | 0 | 0 | 21 | 11 | **41** |
| *Tricalysia* | 0 | 1 | 0 | 0 | 0 | 0 | 7 | 2 | **10** |
| *Vaccinium* | 0 | 9 | 0 | 20 | 0 | 16 | 40 | 3 | **88** |
| *Viburnum* | 0 | 0 | 0 | 4 | 0 | 0 | 121 | 0 | **125** |
| *Vitex* | 0 | 0 | 0 | 0 | 3 | 0 | 1 | 0 | **4** |
| ***TOTAL*** | **10** | **379** | **30** | **212** | **295** | **124** | **1439** | **500** | **2989** |
